# Bub3 and Bub1 maintain the balance of kinetochore-localized Aurora B Kinase and Protein Phosphatase I to Regulate Chromosome Segregation and Anaphase Onset in Meiosis

**DOI:** 10.1101/778480

**Authors:** Gisela Cairo, Anne M. MacKenzie, Soni Lacefield

## Abstract

Accurate chromosome segregation depends on proper attachment of kinetochores to spindle microtubules prior to anaphase onset. The Ipl1/Aurora B kinase corrects improper attachments by phosphorylating kinetochore components and so releasing aberrant kinetochore-microtubule interactions. The localization of Ipl1 to kinetochores in budding yeast depends upon multiple pathways, including the Bub1/Bub3 pathway. We show here that in meiosis, Bub3 is crucial for correction of attachment errors. Depletion of Bub3 results in reduced levels of kinetochore-localized Ipl1, and concomitant massive chromosome mis-segregation caused by incorrect chromosome-spindle attachments. Depletion of Bub3 also results in shorter metaphase I and metaphase II due to premature localization of protein phosphatase 1 (PP1) to kinetochores, which antagonizes Ipl1-mediated phosphorylation. We propose a new role for the Bub1-Bub3 pathway in maintaining the balance between kinetochore-localization of Ipl1 and PP1, a balance that is essential for accurate meiotic chromosome segregation and timely anaphase onset.

**Summary:** Cairo *et al* show that in *S. cerevisiae* meiosis, spindle checkpoint proteins Bub1 and Bub3 have an essential role in preventing chromosome mis-segregation and setting the normal duration of anaphase I and anaphase II onset by regulating the kinetochore-localization of Ipl1 and PP1.

## Introduction

Faithful chromosome segregation in meiosis relies on proper attachment of chromosomes to spindle microtubules. These attachments form through kinetochores, large protein complexes assembled on centromeres that bind microtubules. In meiosis, DNA replication is followed by two rounds of chromosome segregation for the formation of haploid gametes from a diploid progenitor. In meiosis I, the kinetochores on the two sister chromatids are linked together and assemble one microtubule binding site (Marston and Wassmann, 2017). The co-orientation of sister kinetochores allows the kinetochores of the paired homologous chromosomes to attach to microtubules emanating from opposite spindle poles for segregation in anaphase I. In meiosis II, sister kinetochores no longer co-orient but instead attach to opposite spindle poles for segregation in anaphase II.

The cell has developed mechanisms for correcting errors in kinetochore-microtubule attachments. For example, if kinetochores of homologous chromosomes are attached to microtubules emanating from the same spindle pole in meiosis I, the Ipl1/Aurora B kinase phosphorylates kinetochore proteins to drive release of the attachments (Monje-Casas et al., 2007; Saurin, 2018; Yu and Koshland, 2007). This error-correction mechanism gives the kinetochore another opportunity to make proper attachments. In mitosis and presumably in meiosis, there is a balance between Ipl1/Aurora B kinase activity and protein phosphatase 1 (PP1) activity. Once proper kinetochore-microtubule attachments are made, protein phosphatase 1 (PP1) binds to the kinetochore and counteracts Ipl1/Aurora B by dephosphorylating Aurora B/Ipl1 substrates, stabilizing microtubule-kinetochore attachments (Liu et al., 2010; Meadows et al., 2011; Nijenhuis et al., 2014; Rosenberg et al., 2011; Suzuki et al., 2018).

In both mitosis and meiosis, the spindle checkpoint monitors kinetochore-microtubule attachments and will send a signal to delay anaphase onset if kinetochores are unattached. To initiate spindle checkpoint signaling, either protein kinase Mps1 or Polo phosphorylates kinetochore protein Knl1/Spc105 on MELT motifs (Espeut et al., 2015; London et al., 2012; Primorac et al., 2013; Shepperd et al., 2012; Yamagishi et al., 2012). Spindle checkpoint protein Bub3 binds to the phosphorylated MELT motifs and also binds Bub1. Bub1 can recruit other spindle checkpoint proteins, ultimately resulting in the formation of the mitotic checkpoint complex that inhibits the activity of the anaphase-promoting complex/ cyclosome (APC/C) to delay anaphase onset until correct attachments are made (Saurin, 2018).

In addition to their role in spindle checkpoint signaling, two spindle checkpoint proteins, Bub1 and Bub3, also ensure that chromosomes biorient, or attach to opposite spindle poles. For its role in biorientation in meiosis and mitosis, Bub1 phosphorylates histone H2A, which recruits shugoshin (Indjeian et al., 2005; Kawashima et al., 2010; Kiburz et al., 2008; Marston and Wassmann, 2017). Budding yeast and *Drosophila* have a single shugoshin (Sgo1 and mei-S332, respectively), whereas fission yeast and mammals have two shugoshins (Sgo1 and Sgo2) (Katis et al., 2010; Kerrebrock et al., 1992; Kitajima et al., 2004; Kitajima et al., 2006; Marston et al., 2004; Rabitsch et al., 2004; Salic et al., 2004). Shugoshins are also required for the protection of centromeric cohesion in meiosis I. Shugoshins function by serving as a platform for the binding of effector proteins such as protein phosphatase 2A (PP2A), condensin, and the chromosome passenger complex (CPC), of which Ipl1/Aurora B is a subunit (Marston, 2015).

In mitosis, several other pathways can recruit Ip11/Aurora B to the kinetochore, in addition to the Bub3-Bub1 pathway. For example, the CPC binds histone H3, when phosphorylated by haspin kinases (Edgerton et al., 2016; Kelly et al., 2010; Niedzialkowska et al., 2012; Wang et al., 2010; Yamagishi et al., 2010). And, in budding yeast, two additional pathways are also known to recruit Ipl1 (the budding yeast Aurora B homolog): i) the CPC binds to kinetochore protein Ndc10, a subunit of the Cbf3 centromere-binding complex, and ii) Sli15-Ipl1 binds to the inner kinetochore COMA complex (Cho and Harrison, 2011; Fischbock-Halwachs et al., 2019; Garcia-Rodriguez et al., 2019; Yoon and Carbon, 1999). In budding yeast mitosis, loss of any one of these pathways does not severely disrupt chromosome biorientation, but may cause an increase in aneuploidy. However, loss of Ipl1 kinase severely disrupts chromosome biorientation, such that most chromosomes are pulled to one spindle pole (Tanaka et al., 2002). These findings suggest that the multiple recruitment pathways are somewhat redundant in ensuring that Ipl1 is kinetochore-localized for error correction of aberrant kinetochore-microtubule attachments in mitosis. Whether the recruitment pathways also have redundant roles in meiosis has not been tested.

Here, we test the importance of Ipl1 recruitment through Bub1 and Bub3 in meiosis. We find that the loss of Bub1, Bub3, or Sgo1 causes a shorter metaphase I and metaphase II and massive chromosome mis-segregation, especially in meiosis II. In sharp contrast, loss of Bub1 and Bub3 in mitosis causes a longer metaphase and does not cause massive chromosome mis-segregation (Kim et al., 2017; Kim et al., 2015; Warren et al., 2002; Yang et al., 2015). We show that these phenotypes are due to lower levels of Ipl1 at the kinetochore. Because Ipl1 and PP1 counteract one another, we asked whether the shorter duration to anaphase onset was due to premature PP1 at the kinetochore. Our results demonstrate that attenuation of kinetochore binding of PP1 suppressed the short metaphase phenotype of Bub1 or Bub3-depleted cells. Overall, our results demonstrate that the Ipl1 kinetochore recruitment pathway through Bub1, Bub3, and Sgo1 is essential for meiosis. In addition, we show that the balance between kinetochore-localized Ipl1 and PP1 sets the duration of meiosis.

## Results

### Anaphase I and anaphase II onset occurs prematurely in cells depleted of Bub1, Bub3, and Sgo1

To investigate the role of Bub3 in meiotic progression, we used the anchor-away technique to deplete Bub3 from the nucleus at meiosis entry (Haruki et al., 2008). We chose to use anchor-away instead of deleting *BUB3* because homozygous *bub3Δ* cells can become aneuploid and sporulate poorly ((Warren et al., 2002); Yang and Lacefield, unpublished). By depleting Bub3 from the nucleus just prior to meiotic initiation, we avoid any mitotic defects that arise from loss of Bub3. The cells expressed Bub3 tagged with FRB and the ribosomal protein Rpl13a was tagged with FKBP12. Upon addition of rapamycin, FRB and FKBP12 make a stable interaction, leading to the removal of Bub3 from the nucleus, approximately 30 mins after rapamycin addition (Figure S1A-B). Two assays verified that the tag did not disrupt Bub3 function in the absence of rapamycin but did give a phenotype similar to *bub3Δ* cells in the presence of rapamycin. First, cells expressing Bub3-FRB grew similarly to wildtype cells on plates containing the microtubule-depolymerizing drug benomyl. In contrast, with addition of rapamycin, cells were sensitive to benomyl, suggesting that they lost mitotic spindle checkpoint activity (Figure S1C). Second, we find that Bub3-FRB cells sporulated normally but when rapamycin was added produced only inviable spores, suggesting that Bub3 was depleted from the nucleus in meiosis (Figure S2A-B). These results are consistent with Bub3-FRB depletion from the nucleus if rapamycin is present. Similarly, because Bub3 forms a complex with Bub1 at the kinetochore, we also used anchor-away to deplete Bub1 from the nucleus.

To monitor the duration of each meiotic stage, cells expressed Spc42, a spindle pole body (SPB) component, tagged with mCherry; Zip1, a synaptonemal complex component tagged with GFP; and, Tub1, α-tubulin that incorporates into microtubules, tagged with GFP (Bullitt et al., 1997; Scherthan et al., 2007; Sym et al., 1993). Although both Zip1 and Tub1 are tagged with GFP, the proteins are temporally and morphologically distinguishable (Tsuchiya et al., 2011; Tsuchiya et al., 2014). The synaptonemal complex assembles and disassembles in prophase I, showing a nuclear haze of Zip1-GFP that disappears before SPB separation and spindle assembly (Carminati and Stearns, 1997; Scherthan et al., 2007; Sym et al., 1993) (Figure 1A). We used time-lapse microscopy, taking images every 10 minutes throughout meiosis. To assess metaphase duration, we measured the time from SPB separation to the initiation of anaphase I spindle elongation and the time from SPB separation in meiosis II to the initation of anaphase II spindle elongation.

**Figure 1.**
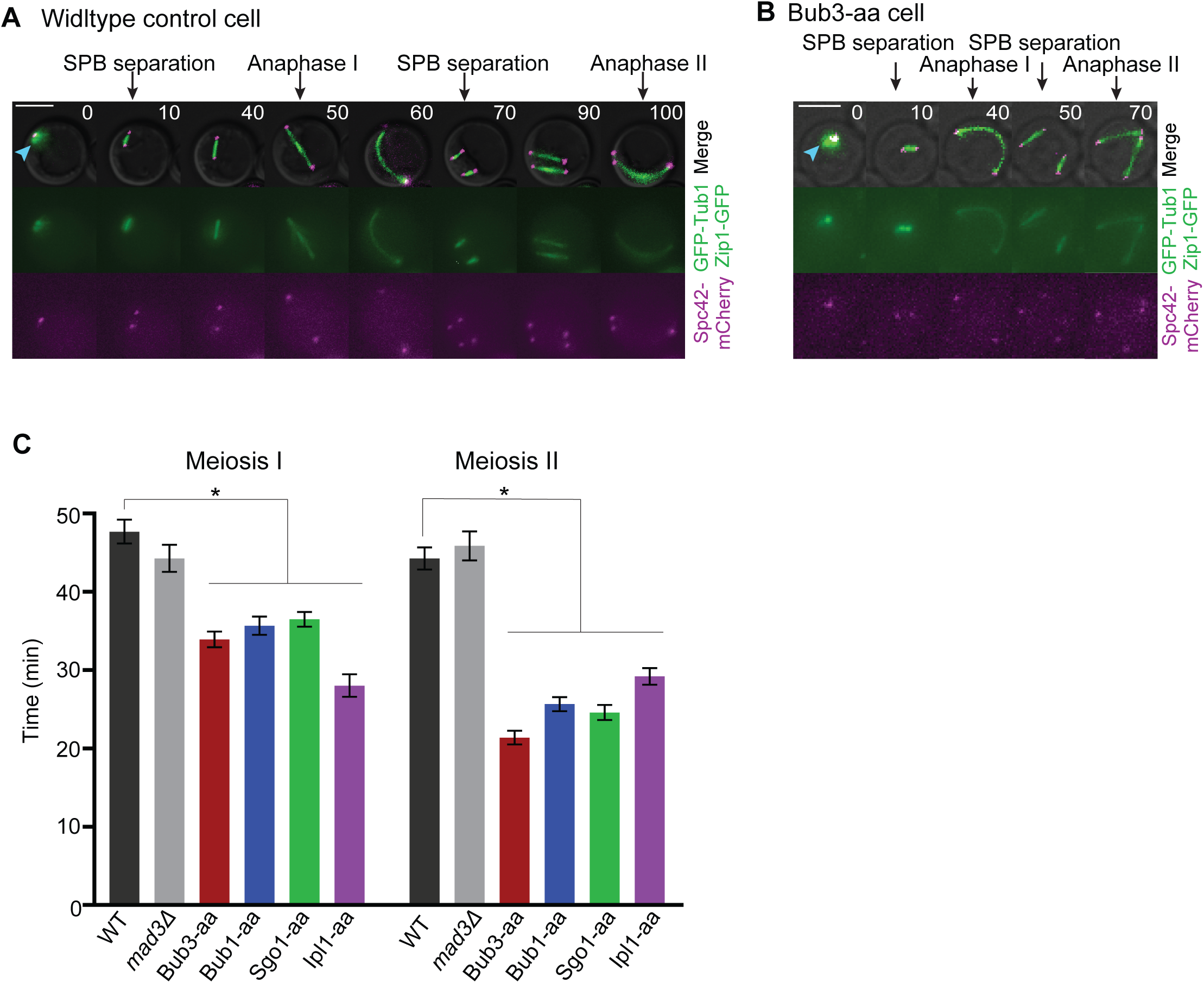
Bub3, Bub1, Sgo1, and Ipl1-depleted cells prevent premature anaphase I and anaphase II onset. (A-B) Representative time-lapse images of (A) a wildtype anchor-away control cell and (B) a cell with Bub3 nuclear depletion (Bub3-aa, anchor away) with rapamycin addition. Cells express Spc42-mCherry, GFP-Tub1, and Zip1-GFP. Blue arrowhead shows Zip1-GFP nuclear localization. Time zero marks prophase I. Scale bars, 5µm. Aa-anchor away. (C) Graph of the mean time from SPB separation to anaphase I onset and SPB separation in meiosis II to anaphase II onset. 100 cells or more from 2 or more independent experiments per genotype were monitored with rapamycin addition. Asterisks indicate a statistically significant difference compared to wildtype anchor-away control cells (p<0.05, Mann-Whitney test). Error bars are the standard error of the mean (SEM).

With the nuclear depletion of Bub3 and Bub1 prior to meiotic initiation, the onset of both anaphase I and anaphase II was shorter in duration than in the wildtype control cells (Figure 1B-C). Anaphase I onset was approximately 14 mins shorter and anaphase II onset was approximately 23 mins shorter in Bub3-depleted cells than in wildtype cells (n =100 cells for each genotype). These results were unexpected because we and others showed that loss of Bub3 or Bub1 in mitosis delays mitotic anaphase onset (Kim et al., 2015; Yang et al., 2015). Therefore, our results suggest that the roles of Bub3 and Bub1 in regulating anaphase onset are different in meiosis than in mitosis.

Although spindle elongation is often used as a marker of anaphase onset, some genetic perturbations can cause premature spindle elongation prior to a true transition into anaphase I or II (Shonn et al., 2000). Therefore, we monitored another marker for anaphase onset, the release of the Cdc14 phosphatase from nucleolus. Cdc14 is normally sequestered in the nucleolus until anaphase I, when it is released; after anaphase I onset, it is re-sequestered, then released again in anaphase II (Marston et al., 2003). We tagged Cdc14 with GFP in cells expressing Spc42-mCherry. In meiosis I, all Bub3-depleted cells released Cdc14-GFP from the nucleolus upon anaphase I spindle elongation, similar to wildtype cells (Figure S3A-C; n= 100 cells for each genotype). In meiosis II, 92% of Bub3-depleted cells released Cdc14 upon spindle elongation. Based on the behavior of the Cdc14 marker, we can conclude that Bub3 is crucial for the normal timing of anaphase onset.

In addition to the activation of spindle checkpoint signaling, the kinetochore-localized Bub3-Bub1 complex is also required for the recruitment of Sgo1 to the kinetochore (Marston, 2015). That localization is important for centromeric cohesion in meiosis I and promotes chromosome biorientation. We asked which role of Bub1 was important for the normal duration of meiosis. Deletion of *MAD3*, which encodes a spindle checkpoint protein that incorporates into the mitotic checkpoint complex for checkpoint signaling, did not result in premature anaphase I or anaphase II onset (Figure 1C; n = 100 cells for each genotype). In contrast, nuclear depletion of Sgo1 through anchor-away resulted in a shorter duration to anaphase I and anaphase II onset compared to the wildtype control strain (Figure 1C). These results suggest that the role of Bub1 in recruiting Sgo1 to the kinetochore is important for setting the normal duration of meiosis.

### Cells depleted of Bub3 and Bub1 undergo massive chromosome mis-segregation in meiosis

We noticed that the spores that formed after meiosis in Bub3 and Bub1-depleted cells often varied in sizes within the same ascus in strong contrast to the production of four spores of equivalent sizes in wild type cells (data not shown). Because spores are packaged mainly around the nucleus, we speculated that the amount of chromatin was different in each spore, perhaps due to massive chromosome mis-segregation, creating some spores with more chromatin than other spores (Neiman, 2011). To monitor the segregation of chromatin, we tagged histone Htb2 with mCherry and performed time-lapse microscopy. We then quantified the areas of the two fluorescent masses after the first meiotic division and the four masses after the second meiotic divisions in wildtype cells and cells depleted of nuclear Bub3 (Figure 2A-D). We found that after meiosis I, 30% of Bub3-depleted cells and 17% of Bub1-depleted cells have uneven chromatin segregation where uneven means greater than one standard deviation from wild type (Figure 2E). The defect is more penetrant in meiosis II; 79% of Bub3-depleted cells and 70% of Bub1-depleted cells have uneven chromatin segregation. Similar to Bub1 depletion, Sgo1 depletion leads to 18% and 71% uneven chromatin segregation in meiosis I and meiosis II, respectively (Figure 2E).

**Figure 2.**
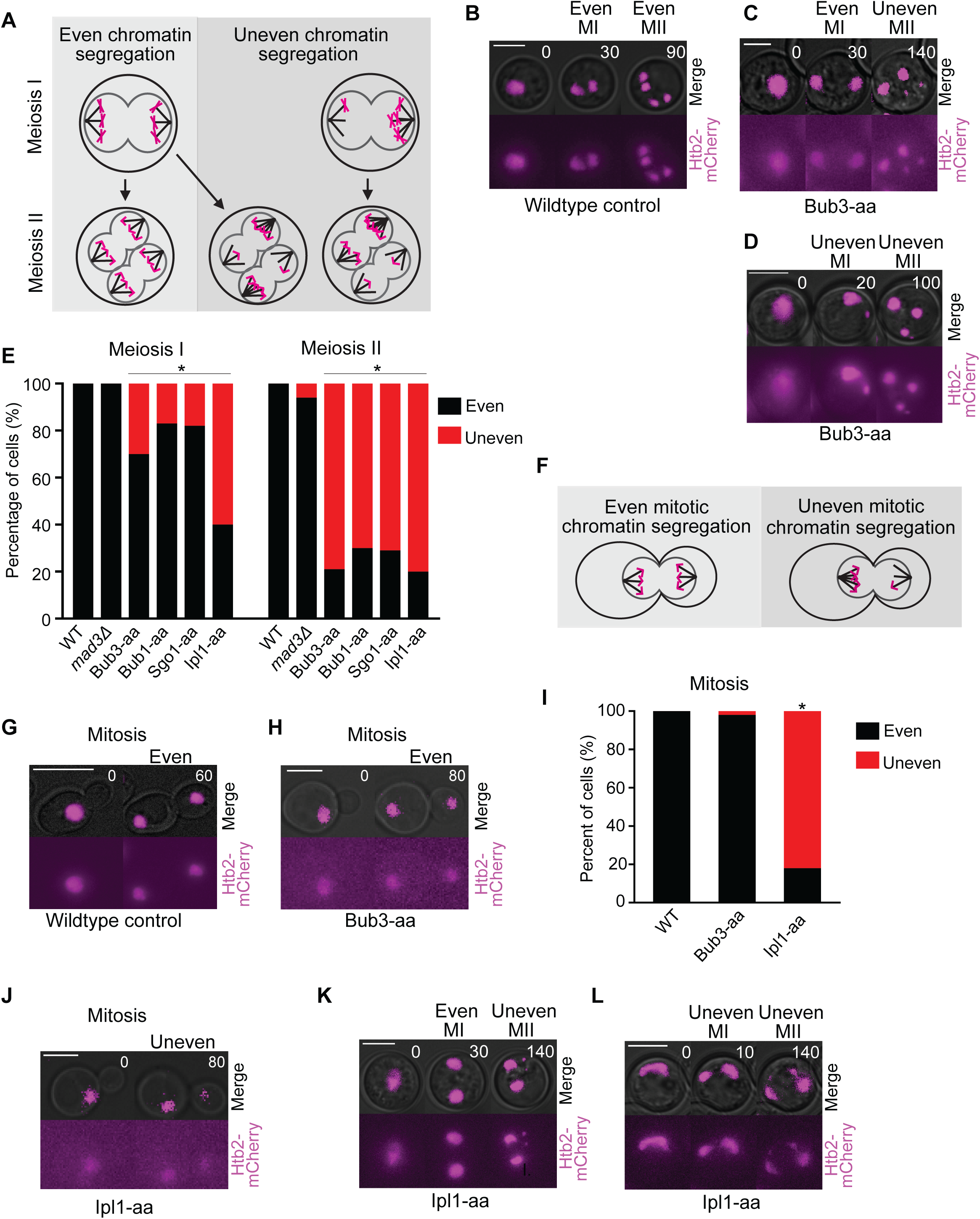
Cells with nuclear depletion of Bub3 display massive chromatin missegregation in meiosis but not in mitosis. (A) Schematic of even and uneven chromatin segregation in meiosis I and meiosis II. (B-D) Representative time-lapses of wildtype cells (B) and cells with nuclear depletion of Bub3 (C-D; Bub3-aa, anchor-away) displaying even and uneven chromatin segregation in meiosis I and meiosis II. Cells express Htb2-mCherry to visualize chromatin. Time zero marks metaphase I. Scale bar, 5µm. (E) Percentage of cells that display even and uneven chromatin segregation in meiosis I and meiosis II. Asterisks indicate a statistically significant difference compared to the wildtype cells (p<0.05, Two-tailed Fisher’s exact test). 100 cells or more from 2 or more independent experiments per genotype were monitored. Anchor-away, aa. (F) Schematic of even and uneven chromatin segregation in a mitotic division. (G-H) Representative time-lapse images of a mitotic division in wildtype cells (G) and cells with nuclear depletion of Bub3 (H; Bub3-aa, anchor-away) showing even chromatin segregation in mitosis. Time zero marks stage before anaphase. Scale bar, 5µm. (I) Percentage of cells that display even and uneven chromatin segregation in mitosis. 100 cells or more from 2 or more independent experiments per genotype. Asterisks indicate a statistically significant difference compared to the wildtype cells (p<0.05, Two-tailed Fisher’s exact test). (J) Representative time-lapse of a mitotic division in cells with nuclear depletion of Ipl1 (Ipl1-aa, anchor-away). (J-K) Representative time-lapses of a meiotic division in cells with nuclear depletion of Ipl1 (H; Ipl1-aa, anchor-away).

The high penetrance of uneven chromatin segregation in cells with nuclear depletion of Bub3 was specific to meiosis. Only 2% of cells showed uneven chromatin segregation in mitosis (Figure 2F-I). Although cells that lack Bub3 have a higher chromosome mis-segregation frequency than wildtype cells in mitosis (Warren et al., 2002), the aneuploidy of a single chromosome is likely not detected with this assay. These results suggest that Bub3 has a more important role in meiotic chromosome segregation than mitotic chromosome segregation.

### Ipl1 levels at the kinetochore are low in Bub3, Bub1, and Sgo1-depleted cells

The uneven chromatin segregation phenotype was reminiscent to that seen in mitosis when cells with a temperature-sensitive mutation of the Aurora B kinase homolog Ipl1 were grown at the restrictive temperature, or in meiosis when Ipl1 was not expressed (Meyer et al., 2013; Tanaka et al., 2002). Ipl1 corrects improper kinetochore-microtubule attachments by phosphorylating kinetochore proteins to release attachments that are not under tension (London and Biggins, 2014). In both mitosis and meiosis, initial chromosome attachments are biased to the old spindle pole body, and Ipl1 is required to release improper attachments for error correction (Meyer et al., 2013; Tanaka et al., 2002). In the absence of Ipl1, chromosomes cannot correct initial attachments, and, subsequently, most chromosomes are pulled towards the old SPB.

We used anchor-away to deplete Ipl1 from the nucleus in cells expressing Htb2-mCherry. We measured chromatin mass size and found that loss of Ipl1 caused an uneven chromatin segregation phenotype in mitosis, meiosis I, and meiosis II (Figure 2E, I-L). In mitosis, 82% of cells with nuclear depletion of Ipl1 displayed an uneven chromatin segregation phenotype (Figure 2I; n=100 cells). In meiosis I, 60% of cells displayed the uneven segregation phenotype, which was more penetrant than the cells with nuclear depletion of Bub3. Interestingly, in meiosis II, 80% of cells in which Ipl1 was depleted from the nucleus displayed an uneven segregation phenotype, which was similar to cells with nuclear depletion of Bub3. These results suggest that the uneven chromatin segregation phenotype in cells depleted of Bub3 and Bub1 is likely due to a loss of error correction, such that most chromosomes segregate to one spindle pole.

To test our hypothesis that kinetochore-localized Ipl1 levels were not sufficient to release aberrant kinetochore-microtubule attachments in meiosis without Bub3, we asked whether chromatin segregation is biased towards the old SPB. We tagged Spc42 with RedStar, a slower maturing red fluorescent protein. When SPBs initially separate, the old SPB will be brighter than the new SPB until RedStar fully matures (Knop et al., 2002; Meyer et al., 2013). We tagged *HTB2* with GFP and assessed whether the larger DNA mass segregated to the old (brighter) SPB or the new (dimmer) SPB (Figure 3A-E). We only scored cells with uneven chromatin segregation. With Bub3, Bub1, and Ipl1 nuclear depletion, the larger DNA mass segregates to the old SPB in more than 88% of cells in meiosis I and meiosis II (Figure 3F; n=100 cells each).

**Figure 3.**
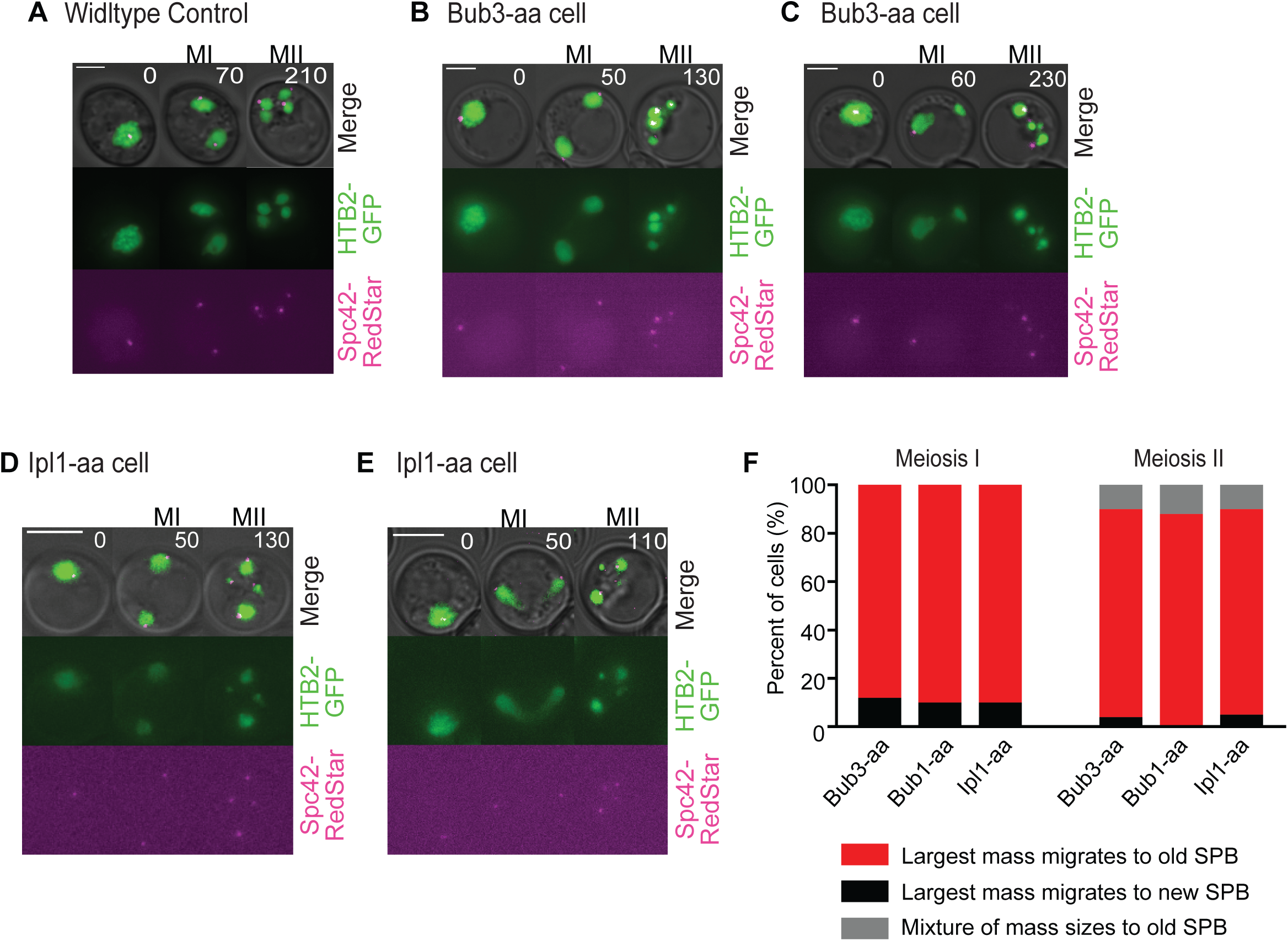
Massive chromatin that missegregates in Bub3-depleted cells mostly migrates to the older SPB, similar to Ipl1-depleted cells. (A-E) Representative time-lapses of wildtype cells (A) and cells with nuclear depletion of Bub3 (B-C) and Ipl1 (D-E) expressing Htb2-GFP and Spc42-RedStar. Time zero marks prophase I. Anchor-away, aa. Scale bars, 5µm. (F) Percentage of cells in which the largest DNA masses migrate to the older and/or brighter SPB for the first and second meiotic division. 100 cells or more from 2 or more independent experiments per genotype were analyzed.

To further test the premise that kinetochores are unable to release improper initial attachments, we monitored a single homologous chromosome pair. We tagged the two homologous chromosome IVs with a LacO array near the centromere in cells expressing GFP-LacI and Spc42-RedStar (Robinett et al., 1996; Straight et al., 1996). We then used 3-minute intervals for time-lapse microscopy and followed the location of the chromosome with respect to the SPBs. We found three different patterns. First, the GFP focus was between the two SPBs, and then properly segregated, suggesting that the chromosomes were bioriented prior to segregation. Second, the GFP focus stayed associated with one SPB and mis-segregated, suggesting that the chromosome was unable to undergo error correction and maintained the initial attachments. Third, the GFP focus traversed back and forth between the SPBs and mis-segregated, suggesting that the chromosome was undergoing error correction, but unable to make proper attachments. In wildtype cells, almost all GFP foci were between the two SPBs and then properly segregate in both meiotic divisions (Figure 4A-B; n=100 cells for each genotype). With Ipl1 nuclear depletion, 96% of the GFP foci stayed attached to one SPB and did not traverse in meiosis I, similar to results from a previous study using a mutant allele of *IPL1* (Meyer et al., 2013). In meiosis II, 95% stayed attached to one SPB, suggesting that initial attachments were unable to correct (Figure 4A, 4C). With Bub3 nuclear depletion, 72% of GFP foci are between the two SPBs and segregate in meiosis I; 24% stayed attached to one SPB and mis-segregated; and, only 4% traversed and mis-segregated. In meiosis II, 72% of GFP foci stayed attached to one SPB (Figure 4A, D-E). These results support the conclusion that with Bub3 nuclear depletion, error correction mechanisms, which release improper kinetochore-microtubule attachments, are not properly functioning in a fraction of cells in meiosis I and the majority of cells in meiosis II. But, in both phases of meiosis, Ipl1 nuclear depletion has a more penetrant defect than Bub3 nuclear depletion, suggesting that a low level of Ipl1 is active at the kinetochore in the absence of Bub3.

**Figure 4.**
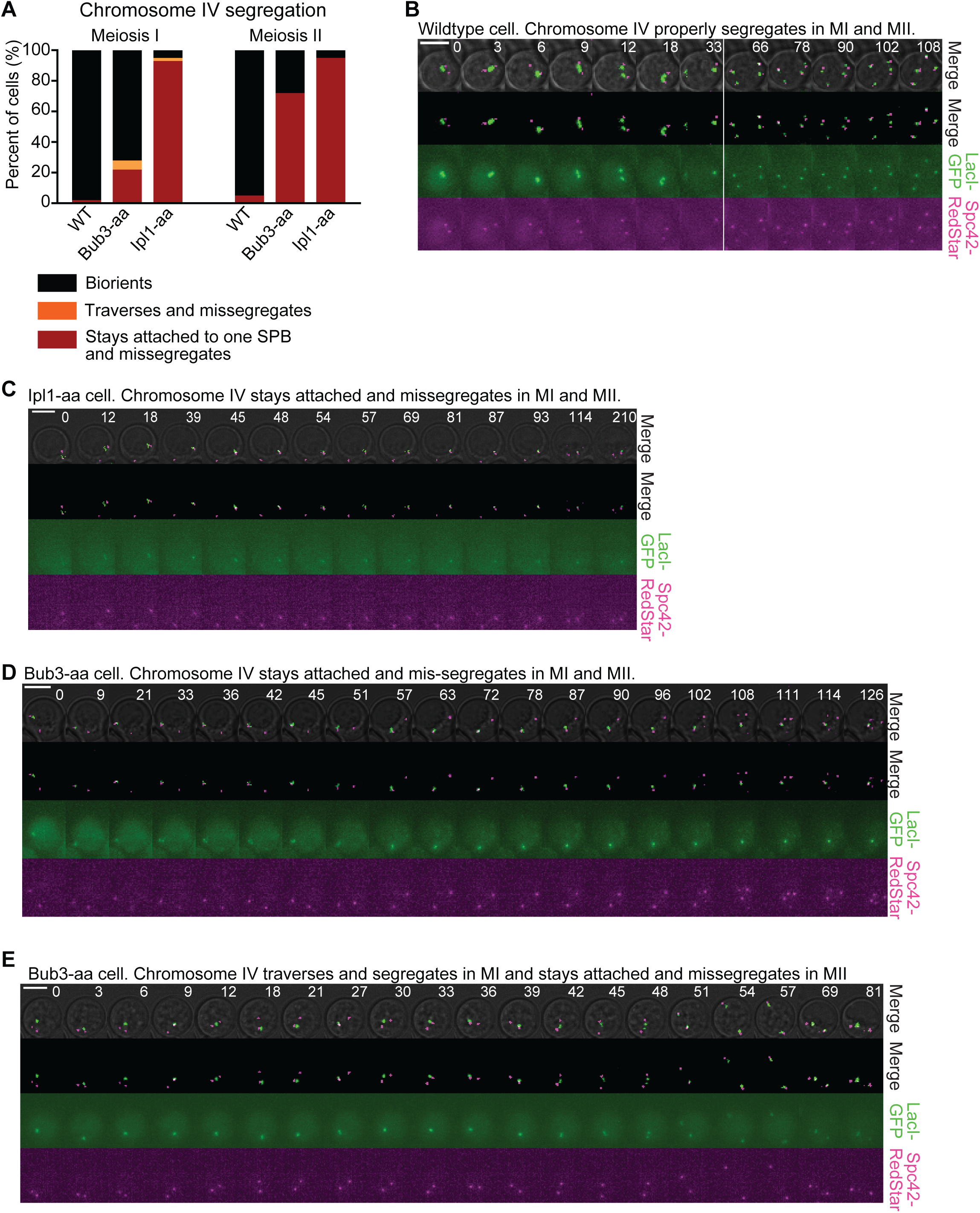
With nuclear depletion of Bub3, chromosome IV usually stays attached to one SPB when it mis-segregates. (A) Percentage of cells classified in the indicated categories according to chromosome IV segregation behavior in the first and second meiotic divisions. 100 cells or more from 3 or more independent experiments per genotype were analyzed. Anchor-away, aa. (B-E) Representative time-lapses of a wildtype cell (B), a cell with nuclear depletion of Ipl1 (C), or Bub3 (D-E). These cells contain LacO repeats near the centromere of chromosome IV and express LacI-GFP and Spc42-RedStar. Time zero marks metaphase I. Scale bars, 5µm.

In mitosis, there are four known Ipl1 kinetochore recruitment pathways, with one of the pathways through Bub3/Bub1/Sgo1(Cho and Harrison, 2011; Edgerton et al., 2016; Fischbock-Halwachs et al., 2019; Garcia-Rodriguez et al., 2019; Kawashima et al., 2010; Peplowska et al., 2014; Verzijlbergen et al., 2014; Yoon and Carbon, 1999). We hypothesize that the Bub3/Bub1/Sgo1 recruitment pathway is crucial for full Ipl1 localization or maintenance in meiosis II. In support of this hypothesis, previous work showed less concentrated levels of Ipl1 at centromeres in the absence of Bub1 in meiosis by immunofluorescence (Yu and Koshland, 2007). However, the reduction was not quantified. We measured the levels of Ipl1-GFP that co-localized with the kinetochore protein Mtw1-mRuby2. Because Ipl1-GFP also localizes to spindle microtubules, we measured the levels at telophase I, after anaphase I spindle breakdown and just before the start of meiosis II (Figure 5A). We found that Ipl1 levels were reduced by 87% with Bub3 depletion (Figure 5B). Overall, our results suggested that the Bub3/Bub1/Sgo1 recruitment pathway is important for maintaining Ipl1 kinetochore levels in meiosis II.

**Figure 5.**
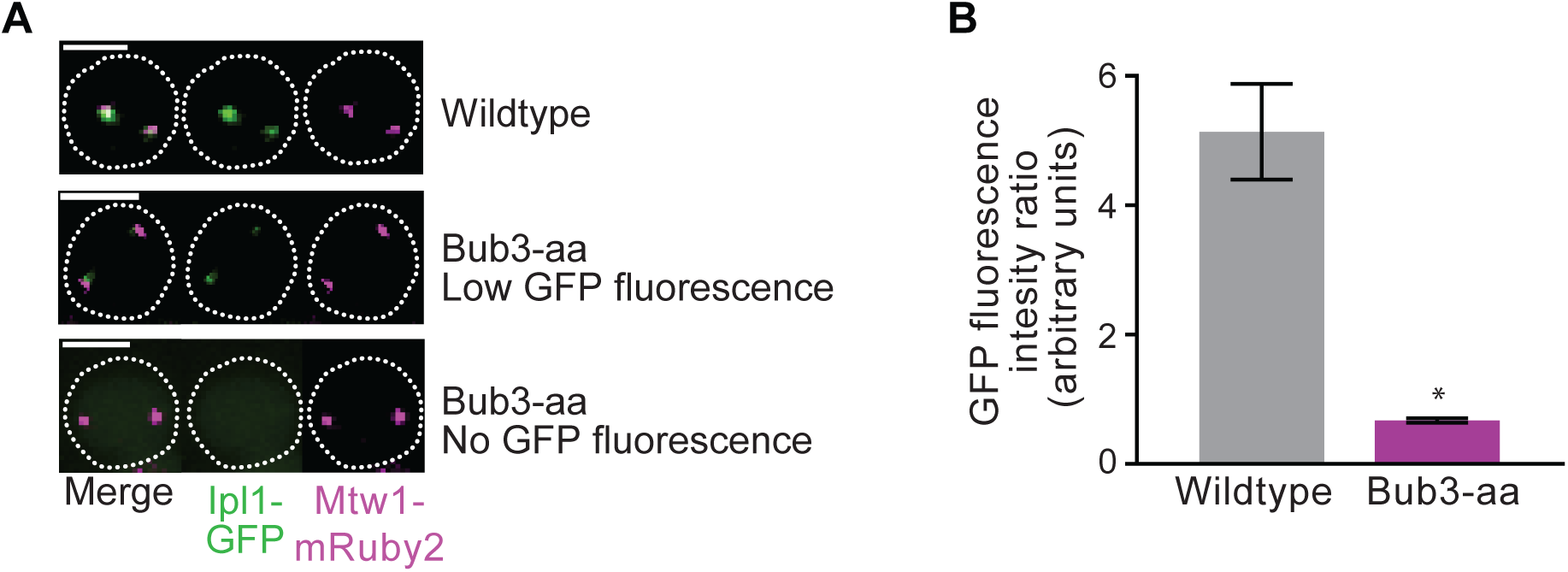
Bub3-depleted cells have reduced kinetochore-localized Ipl1 levels in telophase I. (A) Representative images of wildtype and cells with nuclear depletion of Bub3 at telophase I. Cells express Ipl1-GFP and Mtw1-mRuby2 to visualize the kinetochore. Anchor-away, aa. Bars, 5µm. (B) Graph showing the mean of Ipl1-GFP fluorescence intensity that co-localizes with kinetochore protein Mtw1-mRuby2 in wildtype cells and cells depleted of nuclear Bub3. Values plotted correspond to the ratio between Ipl1-GFP and Mtw1-mRuby2 fluorescence intensity at each Mtw1-mRuby2 spot; fluorescence background levels were subtracted for both channels. Error bars are the SEM. 100 cells or more from 2 or more independent experiments per genotype were monitored. Asterisks indicate a statistically significant difference compared to wildtype cells (*p<0.05, Mann-Whitney test).

### Premature PP1 kinetochore localization advances anaphase I and anaphase II onset

We hypothesized that the reduced levels of Ipl1 at the kinetochore in Bub3, Bub1, and Sgo1-depleted cells could also contribute to premature anaphase onset. To test this hypothesis, we measured the duration from SPB separation to anaphase I or anaphase II onset in cells depleted of Ipl1. We found that both metaphase I and metaphase II were shorter with Ipl1 depletion than in the wildtype control cells, similar to cells depleted of Bub3, Bub1, and Sgo1 (Figure 1C; n= 100 cells).

At the kinetochore, Ipl1 kinase activity and PP1 phosphatase activity counteract one another (Emanuele et al., 2008; Francisco et al., 1994; Hsu et al., 2000; Pinsky et al., 2009). In mammalian cells, Aurora B phosphorylates the PP1 binding site, weakening the interaction between PP1 and KNL1 (Liu et al., 2010). Once kinetochores are bioriented, Aurora B activity at the kinetochore decreases and PP1 binds KNL1 to reverse the phosphorylations on Aurora B substrates. Because PP1 can regulate the normal timing of mitotic progression (Kim et al., 2017; Suzuki et al., 2018), we hypothesized that perhaps reduced Ipl1 kinetochore localization in Bub3 depleted meiotic cells could lead to premature PP1 localization and activity at the kinetochore. To determine if premature PP1 localization leads to shorter metaphase in Bub3 depleted cells, we disrupted a kinetochore binding site for Glc7, the catalytic subunit of PP1. Mutation of the Spc105/KNL1 RVSF motif to RASA disrupts checkpoint silencing, a major function of PP1 at the kinetochore, suggesting that the enzyme is not present for checkpoint silencing (Hendrickx et al., 2009; Liu et al., 2010; Rosenberg et al., 2011). Because, the Spc105^RASA^ mutants are lethal due to a failure to silence the checkpoint, we engineered strains such that wildtype Spc105 was present in mitosis and Spc105^RASA^ was present in meiosis when Bub3 was anchored away. We tagged both *SPC105* and *BUB3* with FRB in our anchor-away strain background and then integrated a copy of *SPC105^RASA^* under the meiosis-specific *REC8* promoter at another locus. Addition of rapamycin depleted both wildtype Spc105 and Bub3, while cells produced Spc105^RASA^ in meiosis. We only analyzed cells that were depleted of Spc105 and Bub3 at prophase I to allow for enough Spc105^RASA^ protein production prior to anchoring away the wildtype protein. As a control, we integrated wildtype *SPC105* under the *REC8* promoter and performed the same analysis.

We found that the control cells depleted of Bub3 and Spc105 in prophase I but expressing the wildtype Spc105 have a premature anaphase I and anaphase II onset, similar to the cells depleted of Bub3 (Figure 6A). In cells with Bub3 and Spc105 depleted in prophase I but expressing Spc105^RASA^, anaphase I and anaphase II onset occurred with similar timing to wildtype cells (45± 1 mins in meiosis I and 45 ± 1 mins in meiosis II, compared to 48 ± 2 mins in meiosis I and 44 ± 1 mins in meiosis II in the wildtype control; average ± SEM; n=100 cells for each). These results suggest that preventing premature PP1 kinetochore localization by mutating the PP1 binding site on the Spc105 receptor rescues the normal timing of anaphase I and anaphase II onset.

**Figure 6.**
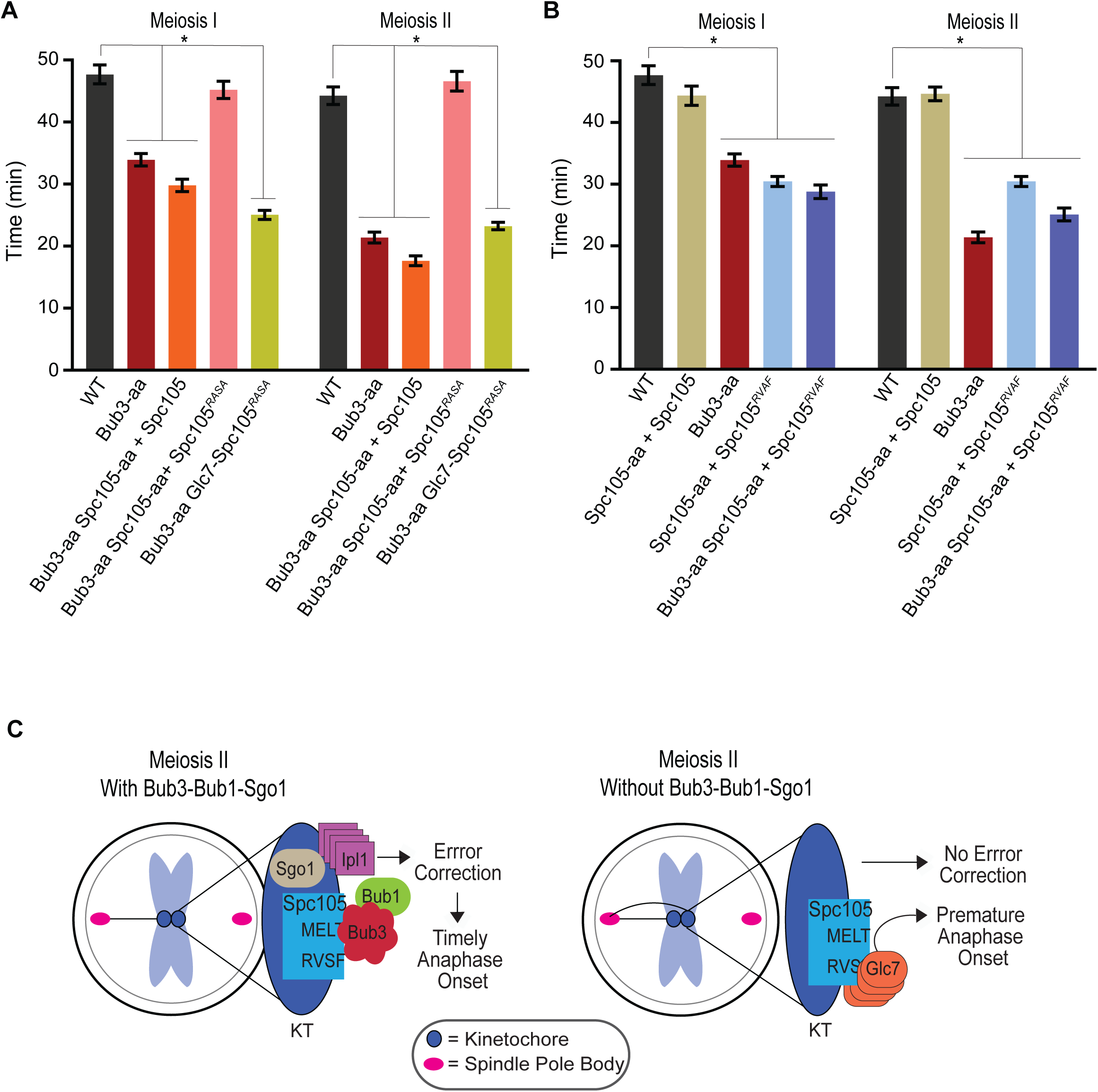
Bub3 regulates the Ipl1/PP1 balance at the kinetochore ensuring proper anaphase onset and chromosome segregation. (A) Graph of wildtype, cells with nuclear depletion of Bub3, and with nuclear depletion of Bub3 and Spc105. Those that have a nuclear depletion of Spc105 also express *SPC105* or *SPC105^RASA^,* under the meiosis-specific *REC8* promoter, at the TRP1 locus. Error bars are the SEM. 100 cells or more from 2 or more independent experiments per genotype were monitored. Asterisks indicate a statistically significant difference compared to wildtype cells (*p<0.05, Mann-Whitney test). Anchor-away, aa. (B) Graph of wildtype, cells with nuclear depletion of Spc105, and cells with nuclear depletion of Bub3. Cells with nuclear depletion of Spc105 express *SPC105* or *SPC105^RVAF^* under the REC8 promoter for meiosis-specific expression, localized at the TRP1 locus. Cells with Glc7-Spc105^RASA^ produce a fusion protein of Glc7 and Spc105 under the control of the *SPC105* promoter at the endogenous locus. Error bars are the SEM. 100 cells or more from 2 or more independent experiments per genotype were monitored. Asterisks indicate a statistically significant difference compared to wildtype cells (*p<0.05, Mann-Whitney test). Anchor-away, aa. (C) Model: Bub3 recruits and maintains Ipl1 at the kinetochore, which ensures proper meiotic chromosome segregation. This prevents PP1 from prematurely binding the kinetochore and premature APC/C activation, which ensures proper anaphase onset. KT, kinetochore.

To further test whether the rescue in timings of Bub3 depleted cells expressing Spc105^RASA^ was due to the loss of PP1 localization, we tethered GLC7 to Spc105^RASA^. In these cells, both anaphase I and anaphase II onset were faster, with similar in timings to cells depleted of Bub3, suggesting that bringing Glc7 back to the kinetochore prevented the rescue of the timings by Spc105^RASA^ (Figure 6A; 25± 1 mins in meiosis I and 23 ± 1 mins in meiosis II, compared to 27 ± 1 mins in meiosis I and 25 ± 1 mins in meiosis II in the Bub3 depleted cells; average ± SEM; n=100 cells for each). Overall, these results support our hypothesis that premature PP1 localization in Bub3 depleted cells results in faster anaphase onset.

Finally, we asked if premature PP1 kinetochore localization, in an otherwise wildtype cell, could also result in faster anaphase onset. We mutated the RVSF motif on Spc105 to RVAF. The Spc105^RVAF^ cannot be phosphorylated by Ipl1 and therefore should allow PP1 to bind prematurely (Hendrickx et al., 2009; Liu et al., 2010; Rosenberg et al., 2011). We anchored-away the wildtype Spc105 and expressed either *SPC105* or *SPC105^RVAF^* from the *REC8* promoter. Expression of *SPC105* showed the normal timing of anaphase I and anaphase II onset (Figure 6B). In contrast, expression of *SPC105^RVAF^* resulted in premature anaphase I and anaphase II onset (Figure 6B; 31± 1 mins in meiosis I and 30 ± 1 mins in meiosis II; average ± SEM; n=100 cells for each). Anchoring away Bub3 did not further decrease the time to anaphase I and anaphase II onset. These results support our hypothesis that premature PP1 localization can indeed lead to faster anaphase onset. Overall, we conclude that cells depleted of Bub3 have less Ipl1 at the kinetochore, leading to premature PP1 localization and faster anaphase onset.

## Discussion

Many mitotic cell-cycle regulators often have modified functions in meiosis important for the meiotic chromosome segregation program. A noteworthy example of a regulatory pathway with increased roles in meiosis is the spindle checkpoint. In addition to delaying anaphase onset if chromosomes are not attached to spindle microtubules, individual spindle checkpoint proteins also have specialized regulatory functions in meiosis. In *C. elegans,* spindle checkpoint proteins MAD-1, MAD-2, and BUB-3 have adopted another function in that they are required for the synapsis checkpoint, which induces apoptosis if cells have not assembled synaptonemal complex during prophase I (Bhalla and Dernburg, 2005; Bohr et al., 2015). The synapsis checkpoint mechanism does not rely on the inhibition of APC/C activity, suggesting that the spindle checkpoint proteins evolved a novel mechanism of action. In another example, the spindle checkpoint protein Mad2 has a more important role in meiosis than in mitosis in regulating chromosome segregation in *S. cerevisiae*. Deletion of *MAD2* in mitosis does not have any major consequence (Li and Murray, 1991). In constrast, in meiosis, *mad2Δ* cells have enhanced chromosome mis-segregation in meiosis I and premature anaphase I onset, likely due to premature APC/C activity (Shonn et al., 2000; Shonn et al., 2003; Tsuchiya et al., 2011). Loss of Mad3 in meiosis does not cause similar phenotypes, although Mad3 is required for spindle checkpoint signaling. These results suggest that Mad2 has an important role independent of spindle checkpoint signaling in regulating meiotic duration and chromosome segregation.

In this study, we show that the spindle checkpoint proteins Bub1 and Bub3 also have more important checkpoint-independent functions in meiosis compared to mitosis. Meiotic nuclear depletion of Bub1 or Bub3 causes a shorter metaphase I and metaphase II. This is in contrast to the results in mitosis in which *bub1Δ* and *bub3Δ* cells have a delayed anaphase onset (Kim et al., 2015; Yang et al., 2015). In addition, we show that Bub1 and Bub3 are crucial in meiosis for regulating bipolar kinetochore-microtubule attachments for proper chromosome segregation. Loss of *BUB1* or *BUB3* in meiosis causes massive chromosome mis-segregation in meiosis II, in which most chromosomes segregate to one spindle pole and less than one percent of the spores are viable. In mitosis, *bub1Δ* and *bub3Δ* cells are viable (Hoyt et al., 1991).

Massive chromosome mis-segregation, in which most chromosomes are pulled to one spindle pole, was a phenotype originally discovered in mitosis with a temperature-sensitive mutation in the Ipl1 kinase (Tanaka et al., 2002). It is now understood that Ipl1 phosphorylates outer kinetochore proteins to release kinetochore-microtubule attachments that are not under tension, allowing another attempt to make proper attachments (London et al., 2012). In addition, Ipl1 is required for spindle checkpoint signaling of tensionless attachments, likely by disrupting kinetochore-microtubule attachments. Therefore, cells that lack Ipl1 cannot correct improper microtubule-kinetochore attachments, and they are unable to signal the spindle checkpoint. In both mitosis and meiosis, the initial attachments of kinetochores mainly occur from microtubules emanating from the old, more mature SPB, such that loss of Ipl1 results in uneven segregation of chromosomes, with the majority of chromosomes traveling to the old SPB (Meyer et al., 2013; Tanaka et al., 2002).

Previous work showed that Bub1 has a role in recruiting Ipl1 to the kinetochore in both mitosis and meiosis (Kawashima et al., 2007; Kawashima et al., 2010; Peplowska et al., 2014; Verzijlbergen et al., 2014; Yu and Koshland, 2007). However, massive chromosome mis-segregation does not occur in the absence of Bub1 and Bub3 in mitosis, likely because there are three other known pathways responsible for bringing Ipl1 to the kinetochore in addition to the Bub3-Bub1 pathway (Cho and Harrison, 2011; Edgerton et al., 2016; Yoon and Carbon, 1999). Interestingly, we find that with the depletion of Bub1 or Bub3, massive chromosome mis-segregation was more penetrant in meiosis II than in meiosis I. We note that although there were fewer cells with massive chromosome mis-segregation, aneuploidy still occurred. Tagging an individual chromosome IV showed that Bub3-depleted cells mis-segregated this chromosome in 28% of meiosis I divisions, similar to previous reports (Marston et al., 2004). These results suggest that although there is enough Ipl1 for most cells to undergo initial rounds of error correction in meiosis I, there is not enough Ipl1 to prevent all chromosome mis-segregation. In meiosis II, our results suggest that when Bub1 or Bub3 is depleted, most cells are unable to undergo even the initial rounds of error correction, leading to massive chromosome mis-segregation in which most chromosomes travel to the old SPB.

Why is chromosome mis-segregation more penetrant in meiosis II than meiosis I or mitosis when Bub1, Bub3, or Sgo1 is depleted? Previous work in mitosis showed that cells lacking Sgo1 initially recruit Ipl1 to the kinetochore but fail to maintain that recruitment in late metaphase (Peplowska et al., 2014). Mitotic cells lacking Bub1 or Bub3 likely have enough Ipl1 at the kinetochore to undergo multiple rounds of error correction before the levels of Ipl1 decrease, preventing massive chromosome mis-segregation. Meiotic cells lacking Bub1 or Bub3 likely have sufficient levels of Ipl1 for a couple of rounds of error correction during metaphase I. However, as cells progress, the kinetochore bound levels of Ipl1 decreases and may not be sufficient for error correction in meiosis II. In support of this conclusion, we find that cells depleted of Bub3 have substantially reduced Ipl1 levels at the kinetochore at telophase I compared to the wildtype control cells. In addition, metaphase II is shorter in the Bub1 or Bub3-depleted cells which may prevent sufficient time for error correction of kinetochores with low levels of Ipl1. Therefore, we propose that maintenance of Ipl1 or continual recruitment of Ipl1 through the Bub3-Bub1 pathway is essential for meiosis II chromosome segregation.

Our findings that depletion of Bub1, Bub3, and Sgo1 resulted in a shorter duration of metaphase I and metaphase II onset were surprising. We previously showed that loss of Bub1 and Bub3 in mitosis displayed the opposite phenotype, a delayed anaphase onset (Yang et al., 2015). However, meiotic cells had a substantially lower level of kinetochore-localized Ipl1, leading us to test whether disruptions in the balance between Ipl1 kinase and its counteracting phosphatase PP1 led to a shorter metaphase. We found that cells depleted of Ipl1 also had a shorter duration to anaphase onset. Furthermore, disruption of the PP1 binding site on kinetochore protein Spc105 rescued the shorter metaphase phenotype of Bub3-depleted cells. And, tethering the PP1 catalytic subunit to Spc105^RASA^ disrupted the rescue. These results suggest that the balance between Ipl1 and PP1 at the kinetochore sets the duration of meiosis such that premature PP1 binding leads to a shorter metaphase I and metaphase II onset. Understanding which Ipl1 and PP1 substrates regulate meiotic progression is an important future direction. Our results support the model that recruitment of Ipl1 through Bub3-Bub1 is essential for maintaining Ipl1 levels for error correction and to prevent PP1 kinetochore-localization until chromosomes have properly attached to spindle microtubules (Figure 6C).

## Materials and methods

### Strains and manipulations

Strains used in this study are derivatives of *Saccharomyces cerevisiae* W303 (Table S1). Gene tagging and deletions were performed using standard PCR-based transformations {Janke, 2004 #277}{Longtine, 1998 #274}. Any manipulations were verified by PCR. The anchor-away strains were built in strain backgrounds previously described but with *BUB3*, *BUB1, SGO1, IPL1, or SPC105* tagged with FRB at the endogenous locus and RPL13A with 2XFKBP12 {Haruki, 2008 #311}. The Spc105^RASA^ and Spc105^RVAF^ strains were made by first cloning *SPC105* under the *REC8* promoter into a yeast integrating plasmid, using site-directed mutagenesis to mutate the RVSF motif and then integrating the plasmid into the *TRP1* locus. The Glc7-Spc105^RASA^ strain was engineered previously and crossed into the anchor-away strains {Rosenberg, 2011 #336}.

### Growth conditions

For all the meiotic experiments, cells were grown overnight in YPD (1% bacto-yeast extract, 2% bacto-peptone, 2% glucose) at 30°C, transferred with a 1:40 dilution to YPA (1% yeast extract, 2% bactopeptone, 1% potassium acetate) for 12-16 hours at 30°C, washed twice with water and then incubated in 1% potassium acetate at 25°C for 7-8 hours. In all experiments with nuclear depletion, except for cells wherein Bub3 was depleted at prophase I (as mentioned in text), rapamycin (1mg/mL) was added at the same time cells were transferred to potassium acetate. To deplete Bub3 at prophase I, rapamycin (1mg/mL) was added after 7 hours of transferring the cells to potassium acetate; only cells in prophase I at the time of rapamycin addition were analyzed. For mitosis experiments, cells were grown in 2X synthetic complete (SC) medium (0.67% bacto–yeast nitrogen base without amino acids, 0.2% dropout mix with all amino acids, and 2% glucose) overnight at 30°C and then diluted and grown to log phase for assessment.

### Microscope image acquisition and time-lapse microscopy

All movies were performed in a chamber mounted on a coverslip coated with Concanavalin A (Sigma). In all meiosis movies, after 6-8 hours of transferring cells to potassium acetate, cells were concentrated and loaded into the coverslip in a chamber. An agar pad containing potassium acetate was used to create a monolayer of cells and removed before imaging; pre-conditioned potassium acetate medium was added. In all mitosis movies, cells were grown overnight in 2XSC medium (0.67% bacto–yeast nitrogen base without amino acids, 0.2% dropout mix with all amino acids, and 2% glucose); rapamycin (1mg/mL) was added to log phase cells before 2-3 hours of imaging.

Cells were imaged at room temperature using a Nikon Ti-E inverted-objective microscope equipped with a 60X oil objective (Plan Apochromat NA 1.4 oil), a Lambda 10-3 optical filter changer and Smartshutter, GFP and mCherry filters (Chroma Technology), and a charge-coupled device camera (CoolSNAP HQ2; Photometrics). During time-lapse imaging, Z stacks of 5 sections of 1.2mm each were acquired in 10 min intervals with exposure times of 60-70ms for brightfield and 700–900ms for GFP and mCherry with neutral density filters transmitting 2–32% of light intensity. While meiosis movies were 12-13 hours long, mitosis movies were 6-8 hours long. Cells expressing Ipl1-GFP MTW1-mRuby2 were mounted (without fixation) on a coverslip and imaged every 10 minutes using 15 Z stacks (0.4µm each) after being in potassium acetate for 9-11 hours; cells in telophase I were used for fluorescence intensity measurements.

Cells producing Htb2-mCherry, Spc42-RedStar and Htb2-GFP, LacI-GFP and Spc42-mCherry, or Bub3-FRB-GFP were imaged at room temperature using a DeltaVision pDV microscope (Applied Precision, Ltd.) equipped with a CoolSNAP HQ2/HQ2-ICX285 camera using a 60X oil objective (U-Plan S-Apochromat-N, 1.4 NA). Images were acquired using SoftWoRx software (GE Heathcare). During time-lapse imaging, five z steps (0.8 µm) were acquired every 10 min for 12 hours except for cells expressing LacI-GFP, wherein images were acquired every 3 minutes for 10-12 hours. The exposure times used to image brightfield, Htb2-mCherry, LacI-GFP, Spc42-mCherry, and Bub3-FRB-GFP were 0.3-0.4ms, 0.02-0.03ms, 0.3-0.4ms, and 0.4ms, 0.3-0.5ms respectively, with neutral density filters transmitting 5-32% of light intensity.

### Image processing

We processed all images using ImageJ (National Institutes of Health). Z-stacks were combined into a single maximum intensity projection with NIS-Elements software (Nikon) to obtain the final images presented in this article. Brightness and contrast were adjusted on entire images only.

### Quantification analysis

Images from cells expressing Htb2-mCherry were quantified using Imarisx64 Software. We measured the area (µm^2^) of each DNA mass after the first and second meiotic division by adding all the area values in each Z stack. To determine even and uneven DNA masses, we measured the DNA masses area after mitosis and the first and second meiotic division of our wildtype and anchor-away strains using Imarisx64. For each wildtype cell, after the mitotic or meiosis I division, the two DNA mass sizes were obtained and then subtracted from one another to get a mass size difference. For meiosis II cells, similar calculations were done with the four DNA mass sizes. The resulting difference values were averaged and the standard deviation was used to calculate a range of what we called ‘even’ DNA masses (average±SD). The same measurements and calculations were performed in the individual cells of the anchor-away strains; anything outside of the ‘even’ range (of average±SD) set by the wildtype cells was considered ‘uneven’. These values were plotted using Prism (GraphPad Software).

### Fluorescence Intensity

To measure Ipl1-GFP fluorescence intensity at the kinetochore (using co-localization with kinetochore protein MTW1-mRuby2) of telophase I cells using ImageJ (FIJI) software, a circle was drawn around MTW1-mRuby2 region and the sum intensity of the mRuby2 channel (M) and GFP channel (I) throughout the Z stacks were recorded. For background subtraction, the circle was moved to several regions not containing MTW1-mRuby2 nor Ipl1-GFP and the sum intensity through the Z stacks was recorded for both mRuby2 (M_background_) and GFP (I_background_); the average of the background fluorescence for each channel was calculated and subtracted from each value above. To determine the fluorescence intensity of Ipl1-GFP at the kinetochore, we calculated the ratio between the Ipl1-GFP and MTW1-mRuby2 values; (I- I_background_)/(M- M_background_). All ratio values were averaged together and plotted using Prism (GraphPad Software).

### Statistical analysis

Statistical analysis was performed using Prism (Graphpad Software). Statistical analysis for anaphase onset timing, fluorescence intensity, and the Bub3-FRB-GFP experiment was done using an unpaired, nonparametric Mann-Whitney test with computation of two-tailed exact p values. To analyze Cdc14 release and DNA masses (HTB2-mCherry strains) percentages, we used the two-sided Fischer’s exact test. The number of data points (n) is indicated in the figure legends. Statistically significant differences (p value was <0.05) is indicated with an asterisk.

### Spotting assay

Spot-test series were 10-fold serial dilutions from saturated overnight cultures. Dilution series were spotted in YPD (1% bacto-yeast extract, 2% bacto-peptone, 2% glucose), YPD + rapamycin (1µg/mL), YPD + benomyl (15µg/mL), and YPD + rapamycin (1µg/mL) + benomyl (15µg/mL). The plates were incubated at 30°C for 36-40 hours before imaging.

## Supplemental material

The supplemental information includes three supplemental figures and one table.

## Acknowledgements

We thank Andreas Hochwagen and Fred Cross for strains. We thank Frank Solomon and members of the Lacefield lab for insightful comments on the manuscript. This work was supported by a grant from the NIH (GM105755).

---

**Figure S1.**
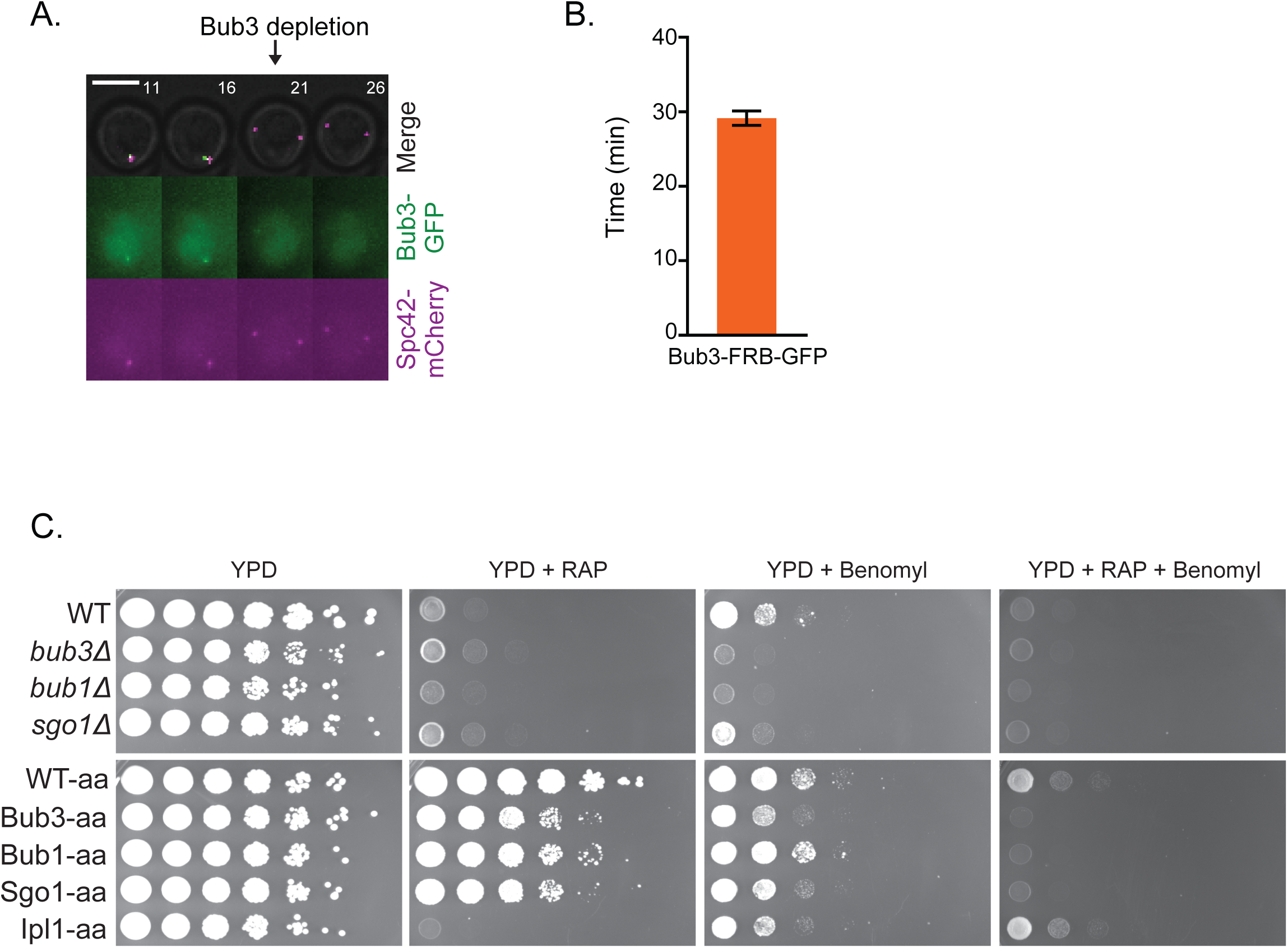
Bub3 is depleted from the nucleus upon rapamycin addition. (A) Representative time-lapse of Bub3-FRB-GFP cell wherein rapamycin is added at t= 0 mins to visualize when Bub3 is anchored away from the nucleus. Bub3-FRB is tagged with GFP and Spc42 is tagged with mCherry. (B). Graph of the mean time of Bub3 nuclear depletion after rapamycin addition. 100 cells from 2 independent experiments were monitored. (C) Mitotic growth of different strains with and without rapamycin and benomyl addition. Cultures are serially diluted 1:10 from a saturated yeast culture. Rapamycin does not affect the mitotic growth of the anchor-away strains due to their genetic background (*tor1-1* mutation). However, the strains without the *tor1-1* mutation are sensitive to rapamycin (wildtype, *bub3Δ, bub1Δ,* and *sgo1Δ*). Final concentrations of benomyl and rapamycin were 15µg/mL and 1µg/mL, respectively. The plates were incubated at 30°C for 36-40 hours before imaging. WT, wildtype. Aa, anchor-away.

**Figure S2.**
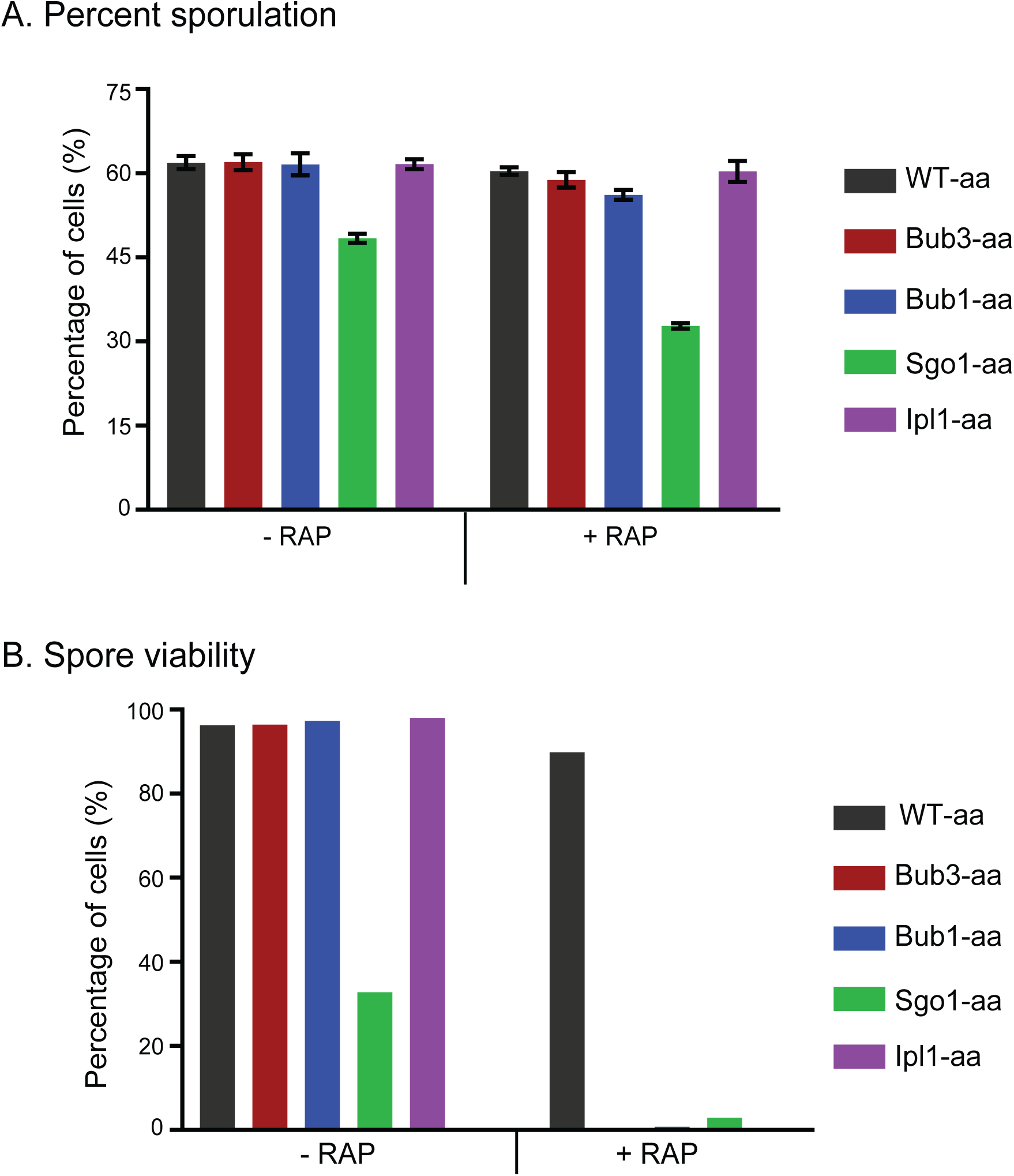
Anchor-away of Bub3, Bub1, Sgo1, and Ipl1 upon rapamycin addition causes loss of spore viability. (A) Percent sporulation of Bub3-aa, Bub1-aa, Sgo1-aa, and Ipl1-aa with the presence and absence of rapamycin. Data obtained from 3 independent experiments. Bars are the SEM. (B) Spore viability of wildtype and Bub3, Bub1, Sgo1, and Ipl1-depleted cells with the presence and absence of rapamycin. Data obtained from analyzing the growth of 240 or more spores for each genotype. WT, wildtype. Aa, anchor-away.

**Figure S3.**
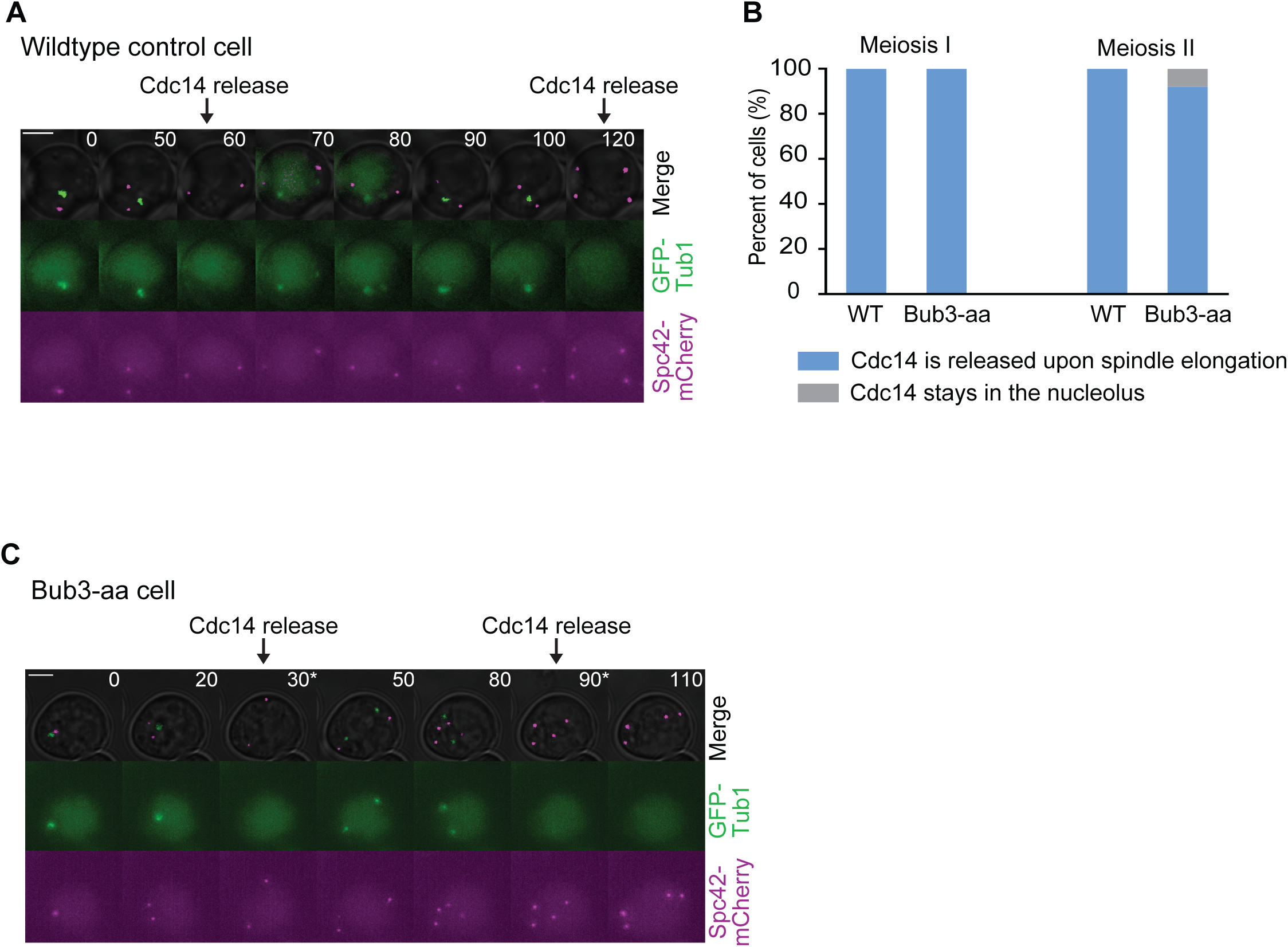
Premature spindle elongation in cells with nuclear depletion of Bub3 coincides with Cdc14-GFP nucleolar release. (A) Representative time-lapse images of a wildtype anchor-away control cell with rapamycin addition. Cell expresses Cdc14-GFP and Spc42-mCherry. Time zero represents prophase I. Scale bars, 5µm. (B) Percentage of wildtype and cells with nuclear depletion of Bub3 wherein Cdc14 was either released or kept in the nucleolus upon spindle elongation. (C) Representative time-lapse images a cell with nuclear depletion of Bub3. Cell expresses Cdc14-GFP and Spc42-mCherry. Time zero represents prophase I. Scale bars, 5µm.

**Table S1.**
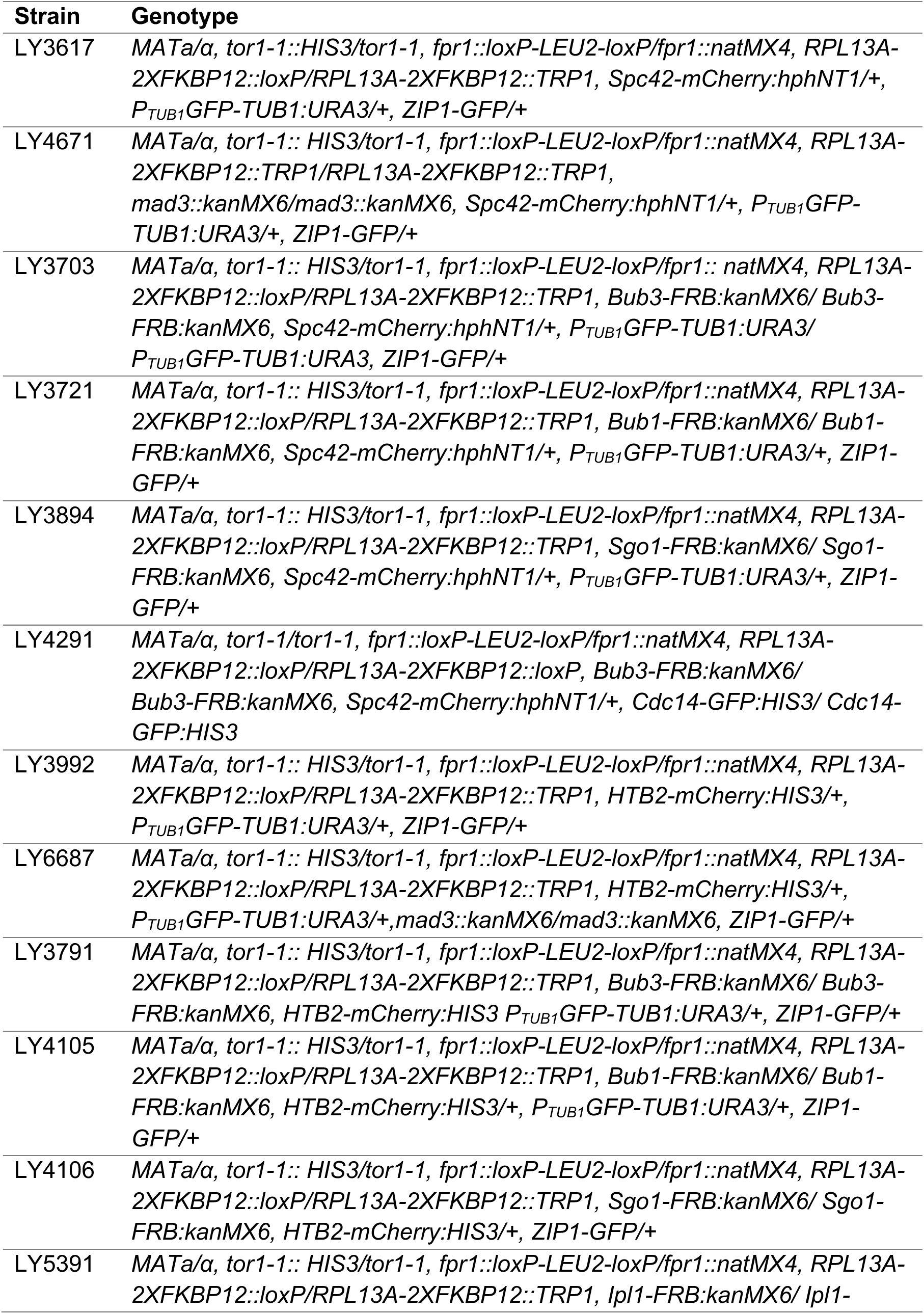
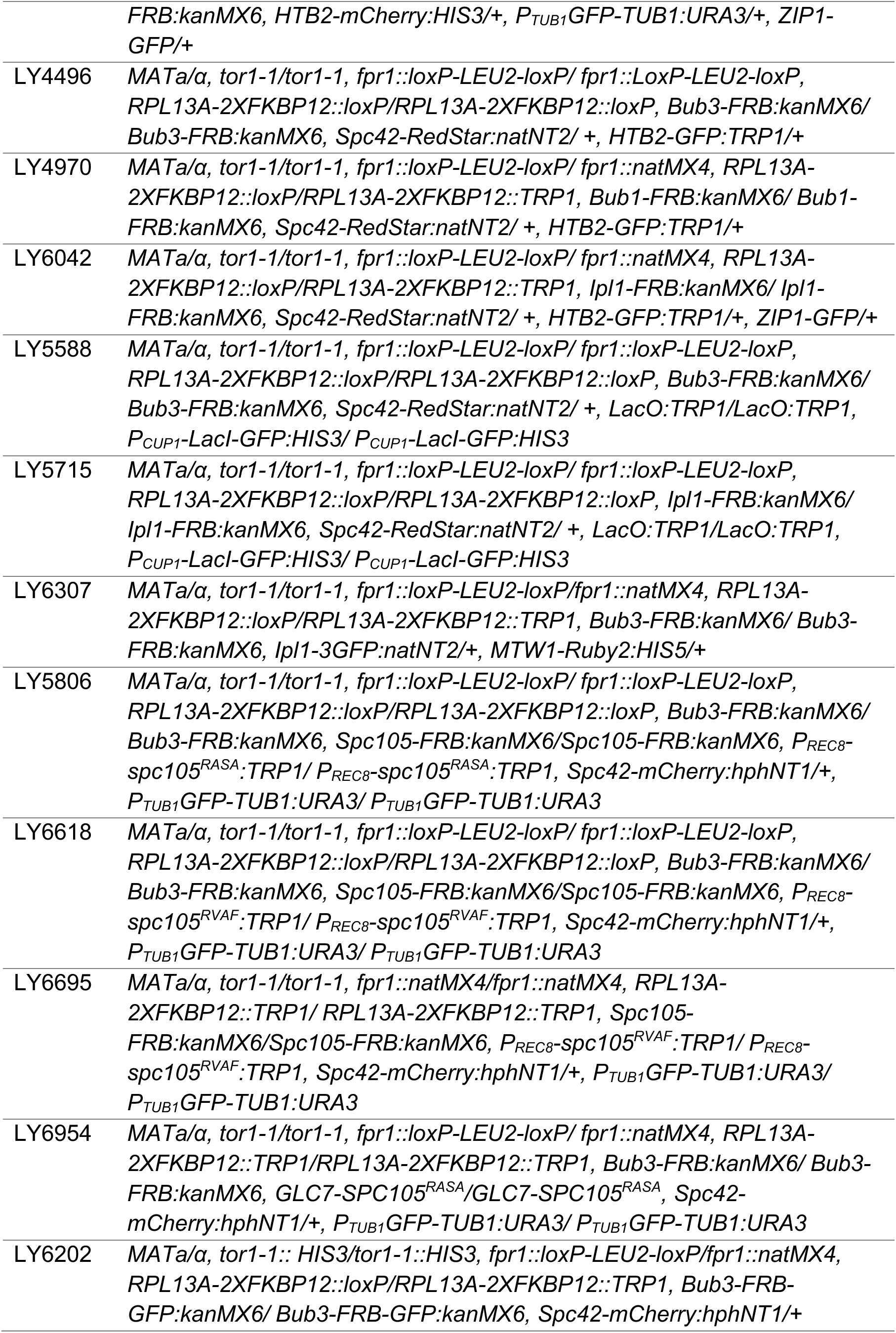
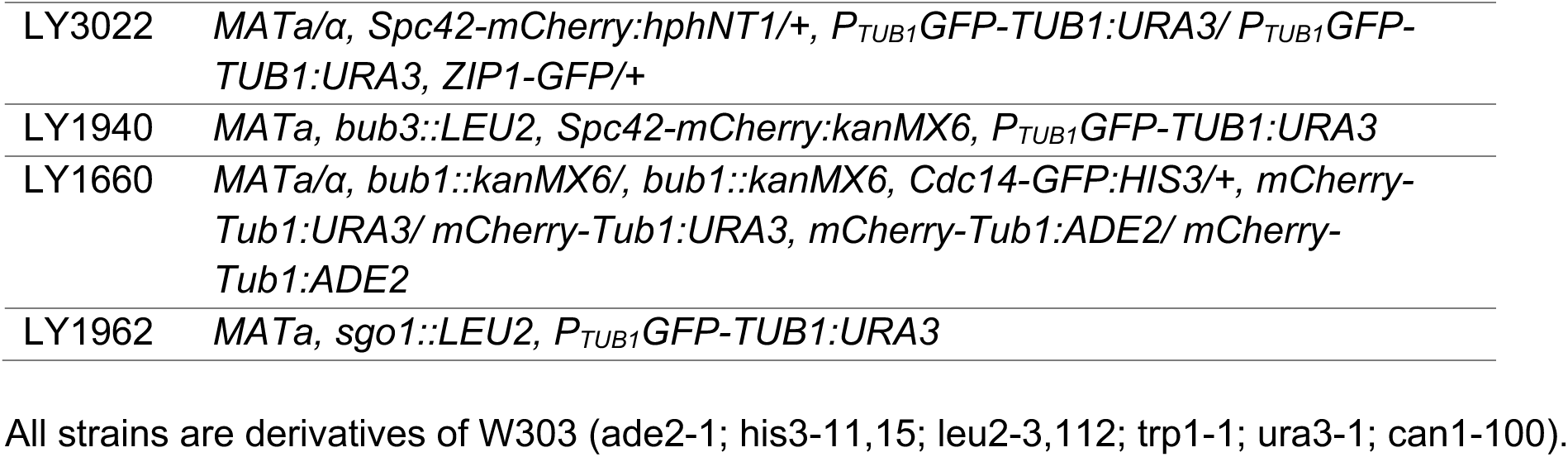
Strains used in this study

| Strain | Genotype |
| --- | --- |
| LY3617 | *MATa/α, tor1-1::HIS3/tor1-1, fpr1::loxP-LEU2-loxP/fpr1::natMX4, RPL13A-2XFKBP12::loxP/RPL13A-2XFKBP12::TRP1, Spc42-mCherry:hphNT1/+, P<sub>TUB1</sub>GFP-TUB1:URA3/+, ZIP1-GFP/+* |
| LY4671 | *MATa/α, tor1-1::HIS3/tor1-1, fpr1::loxP-LEU2-loxP/fpr1::natMX4, RPL13A-2XFKBP12::TRP1/RPL13A-2XFKBP12::TRP1, mad3::kanMX6/mad3::kanMX6, Spc42-mCherry:hphNT1/+, P<sub>TUB1</sub>GFP-TUB1:URA3/+, ZIP1-GFP/+* |
| LY3703 | *MATa/α, tor1-1::HIS3/tor1-1, fpr1::loxP-LEU2-loxP/fpr1::natMX4, RPL13A-2XFKBP12::loxP/RPL13A-2XFKBP12::TRP1, Bub3-FRB:kanMX6/ Bub3-FRB:kanMX6, Spc42-mCherry:hphNT1/+, P<sub>TUB1</sub>GFP-TUB1:URA3/ P<sub>TUB1</sub>GFP-TUB1:URA3, ZIP1-GFP/+* |
| LY3721 | *MATa/α, tor1-1::HIS3/tor1-1, fpr1::loxP-LEU2-loxP/fpr1::natMX4, RPL13A-2XFKBP12::loxP/RPL13A-2XFKBP12::TRP1, Bub1-FRB:kanMX6/ Bub1-FRB:kanMX6, Spc42-mCherry:hphNT1/+, P<sub>TUB1</sub>GFP-TUB1:URA3/+, ZIP1-GFP/+* |
| LY3894 | *MATa/α, tor1-1::HIS3/tor1-1, fpr1::loxP-LEU2-loxP/fpr1::natMX4, RPL13A-2XFKBP12::loxP/RPL13A-2XFKBP12::TRP1, Sgo1-FRB:kanMX6/ Sgo1-FRB:kanMX6, Spc42-mCherry:hphNT1/+, P<sub>TUB1</sub>GFP-TUB1:URA3/+, ZIP1-GFP/+* |
| LY4291 | *MATa/α, tor1-1/tor1-1, fpr1::loxP-LEU2-loxP/fpr1::natMX4, RPL13A-2XFKBP12::loxP/RPL13A-2XFKBP12::loxP, Bub3-FRB:kanMX6/ Bub3-FRB:kanMX6, Spc42-mCherry:hphNT1/+, Cdc14-GFP:HIS3/ Cdc14-GFP:HIS3* |
| LY3992 | *MATa/α, tor1-1::HIS3/tor1-1, fpr1::loxP-LEU2-loxP/fpr1::natMX4, RPL13A-2XFKBP12::loxP/RPL13A-2XFKBP12::TRP1, HTB2-mCherry:HIS3/+, P<sub>TUB1</sub>GFP-TUB1:URA3/+, ZIP1-GFP/+* |
| LY6687 | *MATa/α, tor1-1::HIS3/tor1-1, fpr1::loxP-LEU2-loxP/fpr1::natMX4, RPL13A-2XFKBP12::loxP/RPL13A-2XFKBP12::TRP1, HTB2-mCherry:HIS3/+, P<sub>TUB1</sub>GFP-TUB1:URA3/+, mad3::kanMX6/mad3::kanMX6, ZIP1-GFP/+* |
| LY3791 | *MATa/α, tor1-1::HIS3/tor1-1, fpr1::loxP-LEU2-loxP/fpr1::natMX4, RPL13A-2XFKBP12::loxP/RPL13A-2XFKBP12::TRP1, Bub3-FRB:kanMX6/ Bub3-FRB:kanMX6, HTB2-mCherry:HIS3 P<sub>TUB1</sub>GFP-TUB1:URA3/+, ZIP1-GFP/+* |
| LY4105 | *MATa/α, tor1-1::HIS3/tor1-1, fpr1::loxP-LEU2-loxP/fpr1::natMX4, RPL13A-2XFKBP12::loxP/RPL13A-2XFKBP12::TRP1, Bub1-FRB:kanMX6/ Bub1-FRB:kanMX6, HTB2-mCherry:HIS3/+, P<sub>TUB1</sub>GFP-TUB1:URA3/+, ZIP1-GFP/+* |
| LY4106 | *MATa/α, tor1-1::HIS3/tor1-1, fpr1::loxP-LEU2-loxP/fpr1::natMX4, RPL13A-2XFKBP12::loxP/RPL13A-2XFKBP12::TRP1, Sgo1-FRB:kanMX6/ Sgo1-FRB:kanMX6, HTB2-mCherry:HIS3/+, ZIP1-GFP/+* |
| LY5391 | *MATa/α, tor1-1::HIS3/tor1-1, fpr1::loxP-LEU2-loxP/fpr1::natMX4, RPL13A-2XFKBP12::loxP/RPL13A-2XFKBP12::TRP1, lpl1-FRB:kanMX6/ lpl1-* |
|  | *FRB:kanMX6, HTB2-mCherry:HIS3/+*, *P<sub>TUB1</sub>GFP-TUB1:URA3/+*, *ZIP1-GFP/+* |
| LY4496 | *MATa/α, tor1-1/tor1-1, fpr1::loxP-LEU2-loxP/ fpr1::LoxP-LEU2-loxP*, *RPL13A-2XFKBP12::loxP/RPL13A-2XFKBP12::loxP*, *Bub3-FRB:kanMX6/ Bub3-FRB:kanMX6*, *Spc42-RedStar:natNT2/ +*, *HTB2-GFP:TRP1/+* |
| LY4970 | *MATa/α, tor1-1/tor1-1, fpr1::loxP-LEU2-loxP/ fpr1::natMX4*, *RPL13A-2XFKBP12::loxP/RPL13A-2XFKBP12::TRP1*, *Bub1-FRB:kanMX6/ Bub1-FRB:kanMX6*, *Spc42-RedStar:natNT2/ +*, *HTB2-GFP:TRP1/+* |
| LY6042 | *MATa/α, tor1-1/tor1-1, fpr1::loxP-LEU2-loxP/ fpr1::natMX4*, *RPL13A-2XFKBP12::loxP/RPL13A-2XFKBP12::TRP1*, *lpl1-FRB:kanMX6/ lpl1-FRB:kanMX6*, *Spc42-RedStar:natNT2/ +*, *HTB2-GFP:TRP1/+*, *ZIP1-GFP/+* |
| LY5588 | *MATa/α, tor1-1/tor1-1, fpr1::loxP-LEU2-loxP/ fpr1::loxP-LEU2-loxP*, *RPL13A-2XFKBP12::loxP/RPL13A-2XFKBP12::loxP*, *Bub3-FRB:kanMX6/ Bub3-FRB:kanMX6*, *Spc42-RedStar:natNT2/ +*, *LacO:TRP1/LacO:TRP1*, *P<sub>CUP1</sub>-LacI-GFP:HIS3/ P<sub>CUP1</sub>-LacI-GFP:HIS3* |
| LY5715 | *MATa/α, tor1-1/tor1-1, fpr1::loxP-LEU2-loxP/ fpr1::loxP-LEU2-loxP*, *RPL13A-2XFKBP12::loxP/RPL13A-2XFKBP12::loxP*, *lpl1-FRB:kanMX6/ lpl1-FRB:kanMX6*, *Spc42-RedStar:natNT2/ +*, *LacO:TRP1/LacO:TRP1*, *P<sub>CUP1</sub>-LacI-GFP:HIS3/ P<sub>CUP1</sub>-LacI-GFP:HIS3* |
| LY6307 | *MATa/α, tor1-1/tor1-1, fpr1::loxP-LEU2-loxP/fpr1::natMX4*, *RPL13A-2XFKBP12::loxP/RPL13A-2XFKBP12::TRP1*, *Bub3-FRB:kanMX6/ Bub3-FRB:kanMX6*, *lpl1-3GFP:natNT2/+*, *MTW1-Ruby2:HIS5/+* |
| LY5806 | *MATa/α, tor1-1/tor1-1, fpr1::loxP-LEU2-loxP/ fpr1::loxP-LEU2-loxP*, *RPL13A-2XFKBP12::loxP/RPL13A-2XFKBP12::loxP*, *Bub3-FRB:kanMX6/ Bub3-FRB:kanMX6*, *Spc105-FRB:kanMX6/Spc105-FRB:kanMX6*, *P<sub>REC8</sub>-spc105<sup>RASA</sup>:TRP1/ P<sub>REC8</sub>-spc105<sup>RASA</sup>:TRP1*, *Spc42-mCherry:hphNT1/+*, *P<sub>TUB1</sub>GFP-TUB1:URA3/ P<sub>TUB1</sub>GFP-TUB1:URA3* |
| LY6618 | *MATa/α, tor1-1/tor1-1, fpr1::loxP-LEU2-loxP/ fpr1::loxP-LEU2-loxP*, *RPL13A-2XFKBP12::loxP/RPL13A-2XFKBP12::loxP*, *Bub3-FRB:kanMX6/ Bub3-FRB:kanMX6*, *Spc105-FRB:kanMX6/Spc105-FRB:kanMX6*, *P<sub>REC8</sub>-spc105<sup>RVAF</sup>:TRP1/ P<sub>REC8</sub>-spc105<sup>RVAF</sup>:TRP1*, *Spc42-mCherry:hphNT1/+*, *P<sub>TUB1</sub>GFP-TUB1:URA3/ P<sub>TUB1</sub>GFP-TUB1:URA3* |
| LY6695 | *MATa/α, tor1-1/tor1-1, fpr1::natMX4/fpr1::natMX4*, *RPL13A-2XFKBP12::TRP1/ RPL13A-2XFKBP12::TRP1*, *Spc105-FRB:kanMX6/Spc105-FRB:kanMX6*, *P<sub>REC8</sub>-spc105<sup>RVAF</sup>:TRP1/ P<sub>REC8</sub>-spc105<sup>RVAF</sup>:TRP1*, *Spc42-mCherry:hphNT1/+*, *P<sub>TUB1</sub>GFP-TUB1:URA3/ P<sub>TUB1</sub>GFP-TUB1:URA3* |
| LY6954 | *MATa/α, tor1-1/tor1-1, fpr1::loxP-LEU2-loxP/ fpr1::natMX4*, *RPL13A-2XFKBP12::TRP1/RPL13A-2XFKBP12::TRP1*, *Bub3-FRB:kanMX6/ Bub3-FRB:kanMX6*, *GLC7-SPC105<sup>RASA</sup>/GLC7-SPC105<sup>RASA</sup>*, *Spc42-mCherry:hphNT1/+*, *P<sub>TUB1</sub>GFP-TUB1:URA3/ P<sub>TUB1</sub>GFP-TUB1:URA3* |
| LY6202 | *MATa/α, tor1-1::HIS3/tor1-1::HIS3*, *fpr1::loxP-LEU2-loxP/fpr1::natMX4*, *RPL13A-2XFKBP12::loxP/RPL13A-2XFKBP12::TRP1*, *Bub3-FRB-GFP:kanMX6/ Bub3-FRB-GFP:kanMX6*, *Spc42-mCherry:hphNT1/+* |
| LY3022 | *MATa/α, Spc42-mCherry:hphNT1/+, P<sub>TUB1</sub>GFP-TUB1:URA3/ P<sub>TUB1</sub>GFP-TUB1:URA3, ZIP1-GFP/+* |
| LY1940 | *MATa, bub3::LEU2, Spc42-mCherry:kanMX6, P<sub>TUB1</sub>GFP-TUB1:URA3* |
| LY1660 | *MATa/α, bub1::kanMX6/, bub1::kanMX6, Cdc14-GFP:HIS3/+, mCherry-Tub1:URA3/ mCherry-Tub1:URA3, mCherry-Tub1:ADE2/ mCherry-Tub1:ADE2* |
| LY1962 | *MATa, sgo1::LEU2, P<sub>TUB1</sub>GFP-TUB1:URA3* |

All strains are derivatives of W303 (ade2-1; his3-11,15; leu2-3,112; trp1-1; ura3-1; can1-100).

